# The diadenosine tetraphosphate hydrolase YqeK controls fitness, biofilm formation, staphyloxanthin production and virulence in *Staphylococcus aureus*

**DOI:** 10.64898/2026.09.08.750060

**Authors:** Vu Van Loi, Sarah Przibilla, Fabiana Burchert, Jan Pane-Farre, Tobias Busche, Christian Rückert-Reed, Gert Bange, Haike Antelmann

## Abstract

Diadenosine tetraphosphate (Ap_4_A) is a nucleotide metabolite, which is degraded by the YqeK hydrolase in *Staphylococcus aureus in vitro*. In this study, we analyzed the phenotypes of the Δ*yqeK* mutant under stress, antibiotics, biofilm and macrophage infection conditions to investigate the functions of Ap_4_A in *S. aureus* COL. Using nucleotide metabolomics, we confirmed that Ap_4_A levels are 105-fold higher in the Δ*yqeK* mutant, accompanied by lower adenylate and guanylate nucleotide pools. The Δ*yqeK* mutant showed a delayed growth in LB, TSB and RPMI medium, and decreased survival under lethal oxidative and quinone stress. Transcriptome analysis revealed the upregulation of the *sspABC* operon and the CodY, T-box Met and G-box regulons indicating increased amino acids and GTP biosynthesis, whereas the AgrA, Fur, PurR, and T-box Cys regulons were downregulated in the Δ*yqeK* mutant. These gene expression changes could be restored to WT level in the *yqeK*^+^ complemented strain, resulting also in a DNA damage response as revealed by the induction of LexA regulon members and mobile genetic elements (pathogenicity island SAPI3 and prophage L54a). Moreover, the Δ*yqeK* mutant showed enhanced biofilm formation, higher intracellular iron levels and lower staphyloxanthin levels. Using infection assays, we demonstrated a decreased survival of the Δ*yqeK* mutant inside J774A.1 murine macrophages, supporting a link between Ap_4_A and pathogenicity regulation *via* Agr-controlled virulence factors in *S. aureus*. Future research should be directed to understand how Ap_4_A regulates nucleotide, iron and amino acid metabolism as well as biofilm and virulence phenotypes in *S. aureus*.

**IMPORTANCE:** *S. aureus* is an important human pathogen, which can cause life-threatening infections especially in immunocompromised patients. Due to the prevalence of multidrug resistant strains, the search for new drug targets is an urgent goal. Diadenosine tetraphosphate (Ap_4_A) has been shown to contribute to stress responses, antibiotic resistance, biofilm development and virulence in bacteria. In this work, we showed that AP4A is upregulated upon deletion of *yqeK* encoding the Ap_4_A hydrolase in *S. aureus* COL. Moreover, the Δ*yqeK* mutant was impaired in growth and survival during oxidative stress and after infection of murine macrophages, indicating that Ap_4_A contributes to the host-pathogen interactions in *S. aureus*. Transcriptome analyses revealed alterations of the nucleotide, amino acid and iron metabolism as well as the downregulation of Agr-controlled cytotoxins, contributing to the lower virulence of the Δ*yqeK* mutant. Altogether, our results provide leads for the design of inhibitors against YqeK to combat *S. aureus* infections.

## INTRODUCTION

*Staphylococcus aureus* is an opportunistic pathogen, which colonizes the skin and the nose of about 20-30% of the healthy human population without causing symptoms of infections (1, 2). However, especially immunocompromised and elderly people are at risk for hospital- or community-acquired *S. aureus* infections, ranging from skin and soft tissue infections to fatal pneumonia, septic shock and osteomyelitis (3–5). Further complications arise due to the emergence of antibiotic-resistant strains, such as methicillin-resistant *S. aureus* (MRSA), which is classified as ESKAPE priority pathogen, causing a high health burden and morbidity rate with limited therapy options (6–9).

*S. aureus* strains are known for their high genome diversity and plasticity, which evolved due to horizontal gene transfer of mobile genetic elements (MGE), such as phages and plasmids, that often carry virulence factors and antibiotics resistance determinants (10, 11). The regulatory networks for virulence factor control are highly complex, involving among others the accessory gene regulator (Agr) quorum-sensing system, the SaeRS, SrrAB and ArlRS two-component systems, several MarR/SarA-family transcriptional regulators (SarA, MgrA, Rot), the SigmaB alternative sigma factor, the GTP-sensing CodY repressor controlling amino acid biosynthesis and the iron uptake repressor Fur (10, 12–21). Furthermore, *S. aureus* utilizes nucleotide second messengers, such as cyclic di-AMP (c-di-AMP) and (p)ppGpp to control virulence, adaptation to osmotic, acid and oxidative stress, stationary phase survival, phagosomal escape, antibiotic resistance and cellular iron homeostasis (22–29).

While the cellular targets for c-di-AMP and (p)ppGpp are well-studied in many bacteria, there is limited knowledge on the role of Ap_4_A, which accumulates during heat and oxidative stress in *Escherichia coli* and effects the survival under stress and starvation (30–33). APA4 is synthesized mainly from ATP and aminoacyl-AMP as side reaction by amino acyl tRNA synthetases, albeit the regulatory mechanism is poorly understood (34, 35). Moreover, also other adenylate-generating enzymes can produce Ap_4_A (30, 33). The breakdown of Ap_4_A to either ATP and AMP or to two molecules ADP is catalyzed by Nudix hydrolases present in all domains of life, as well as the phosphatase ApaH in Gram-negative bacteria and the HD domain enzyme YqeK in Gram-positive bacteria (30, 36, 37). Thus, Ap_4_A has been also recognized as damaging metabolite, since it is generated as side product of cellular metabolism at low levels and there were no specific signalling pathways known, which are affected directly by Ap_4_A (38).

However, increasing evidence suggests that Ap_4_A acts as *bona fide* nucleotide second messenger. Recent studies have identified specific cellular targets of Ap_4_A, providing further evidence for its role as a *bona fide* signalling molecule in bacteria. In *Bacillus subtilis*, Ap_4_A binds to inosine 5’-monophosphate dehydrogenase (IMPDH) and promotes formation of a less active oligomeric state, thereby modulating purine nucleotide homeostasis and adaptation to heat stress (39). Moreover, Ap4A binds to the cystathionine-beta-synthase (CBS) domain of AcuB, stabilizing AcuB and enhancing its inhibition of the histone deacetylase (HDAC)-like deacetylase AcuC, thereby linking Ap4A signaling to reversible protein acetylation and acetyl-CoA metabolism (40, 41). Thus, tight control of intracellular Ap4A levels is likely essential for regulating its signaling functions and preventing inappropriate modulation of its cellular targets.

In Gram-negative bacteria, degradation of Ap_4_A has been associated with changes in bacterial physiology and adaptation processes, including heat and oxidative stress resistance, protein homeostasis, antibiotics resistance, cell division, motility and biofilm formation (30, 33, 36). Furthermore, Ap_4_A has been implicated in the regulation of pathogenicity in *Pseudomonas aeruginosa* and *Salmonella* Typhimurium (42, 43). In *P. aeruginosa*, the Δ*apaH* mutant showed decreased survival in lettuce leaves, *Galleria mellonella* and mouse infection models. This virulence phenotype was reproducible in clinical *P. aeruginosa* isolates harboring the Δ*apaH* deletion (42). Transcriptome analyses supported the downregulation of key virulence factors, such as extracellular proteases, elastase, iron-uptake systems and quorum-sensing molecules upon Ap_4_A accumulation in the Δ*apaH* mutant, indicating an important regulatory role of Ap_4_A to modulate diverse virulence traits (42). In *S*. Typhimurium, the absence of ApaH and the Nudix hydrolase YgdP alone and in combination led to increased Ap_4_A levels and impaired adherence and intracellular invasion of mammalian HEp-2 and U-937 cell cultures (43). Furthermore, in *Helicobacter pylori* the NudA hydrolase is constitutively expressed and essential for oxidative stress resistance (44).

The YqeK homologs of *S. aureus*, *B. subtilis*, *Streptococcus pyogenes* and *Streptococcus pneumoniae* have been recently structurally and biochemically characterized, and it was confirmed that the level of Ap_4_A was elevated in the *B. subtilis* Δ*yqeK* mutant, supporting that YqeK functions as Ap_4_A hydrolase to cleave Ap_4_A symmetrically into two ADP molecules *in vitro* (37, 45–47). Recent work also showed that Np_4_-capped mRNAs increase upon Np_4_A accumulation in the *B. subtilis* Δ*yqeK* mutant, indicating that YqeK also functions as decapping enzyme that deprotects 5’ ends of mRNAs to facilitate mRNA degradation of Np_4_-capped mRNAs (47, 48). Additionally, a study conducted in parallel to ours reported a role of YqeK in the nitrosative and acid stress response and transcriptome changes associated with virulence and cellular metabolism in the *S. aureus* USA300 LAC Δ*yqeK* mutant (49).

In our work, we combined metabolomics, transcriptomics and phenotypes analyses to investigate the physiological role of the Ap_4_A hydrolase YqeK in *S. aureus* COL. Nucleotide LC-MS analyses showed significantly increased Ap_4_N levels in the Δ*yqeK* mutant. RNA-seq transcriptome analyses revealed differential expression of regulons involved in amino acid, nucleotide and iron metabolism, as well as the downregulation of the Agr virulence regulon in the Δ*yqeK* mutant. Furthermore, the Δ*yqeK* mutant showed a growth delay, decreased survival under diamide and quinone stress and after infection of J774A.1 murine macrophages, increased biofilm formation and total iron levels, and decreased staphyloxanthin production. Together, these findings link Ap_4_A accumulation to altered metabolism, reduced fitness and virulence in *S. aureus* COL.

## RESULTS

### The *S. aureus* COL Δ*yqeK* mutant shows increased Ap_4_A levels and decreased nucleotide pools in the metabolome

To investigate the role of Ap_4_A in *S. aureus* COL, we constructed a Δ*yqeK* deletion mutant and a *yqeK*^+^ complemented strain. Using liquid chromatography mass spectrometry (LC-MS)-based nucleotide metabolomics, we analyzed the nucleoside polyphosphate and nucleotide pools of the WT, Δ*yqeK* mutant and *yqeK*^+^ strains (**Fig. 1**). The MS results revealed 105-fold higher Ap_4_A levels in the Δ*yqeK* mutant, while the levels of Ap_4_U and Ap_4_G were 6- and 5-fold increased, respectively, compared to the WT (**Fig. 1A**). Comparison of the nucleotide pools revealed a two-fold decrease of ATP, ADP, and AMP in the Δ*yqeK* mutant, consistent with the higher Ap_4_A pool (**Fig. 1B**). Similarly, the levels of UDP, UTP, GMP, GTP, CMP, and CTP were 2-3-fold decreased upon YqeK depletion (**Figs. 1C, D, E**). The complementation with YqeK restored Ap_4_N and the nucleotide pools to WT level. Together, the metabolomics results revealed that YqeK is important for degradation of AP4N and maintenance of nucleotide homeostasis in *S. aureus* COL.

**Fig. 1.**
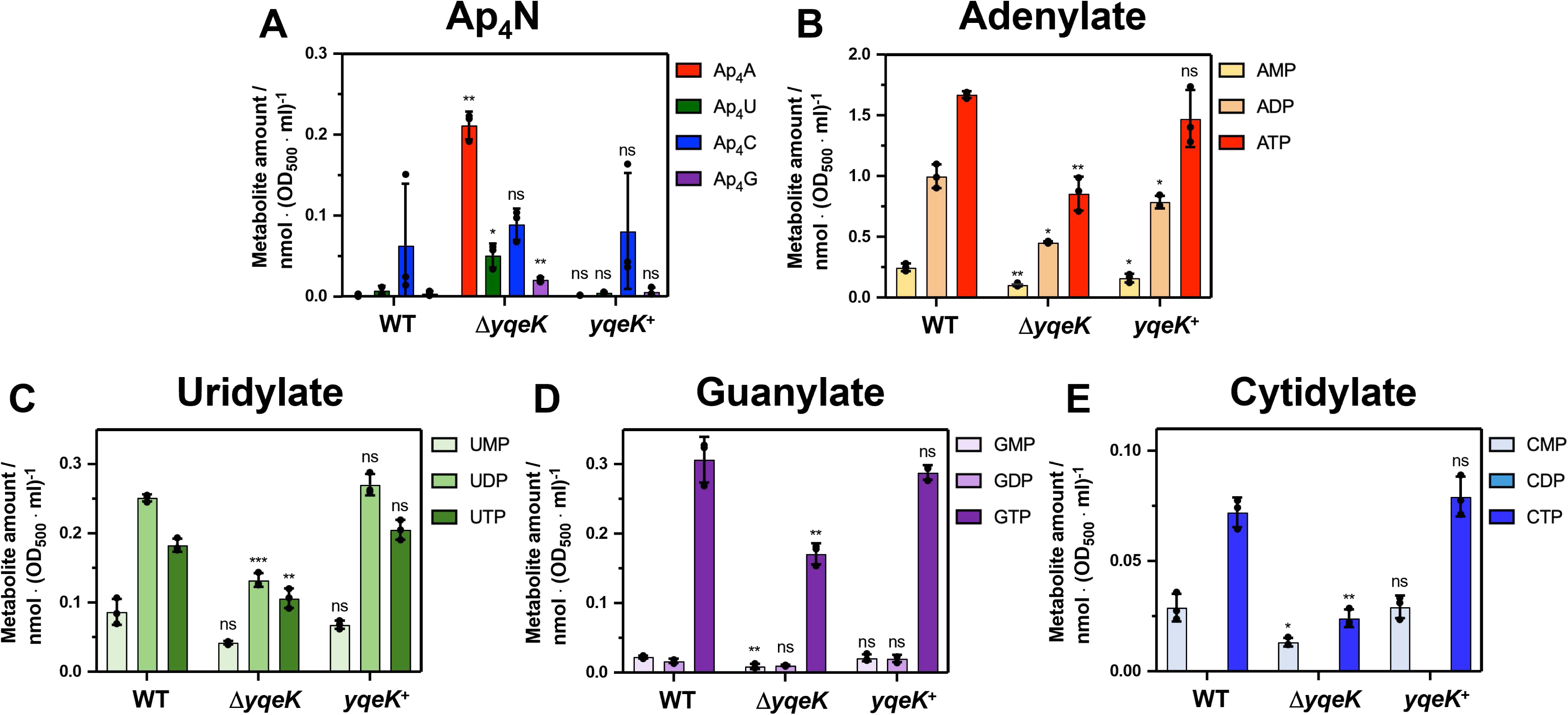
The Δ*yqeK* mutant exhibits increased Ap_4_A levels and decreased nucleotide pools in *S. aureus* COL. *S. aureus* COL WT, Δ*yqeK* mutant and *yqeK*^+^ complemented strains were grown in RPMI with 1% (w/v) glucose and 1% (w/v) xylose in 3 biological replicates with 3 technical replicates each. Cells were harvested at an OD_500_ of 1 and intracellular metabolites were extracted and analyzed by LC-MS as described in the Methods section. The intracellular concentrations of Ap_4_A, Ap_4_U, Ap_4_C and Ap_4_G **(A)**, AMP, ADP and ATP **(B)**, UMP, UDP and UTP **(C)**, GMP, GDP and GTP **(D)**, and CMP, CDP and CTP **(E)** were calculated as nmol per OD_500_ x culture volume. Mean values are presented and error bars represent the standard deviation (SD). Statistical significance was determined using an unpaired two-tailed Student’s *t*-test comparing Δ*yqeK* and *yqeK*^+^ *vs*. the WT (^ns^, *p* > 0.05; *, *p* ≤ 0.05; **, *p* ≤ 0.01; ***, *p* ≤ 0.001).

### The Δ*yqeK* mutant shows significantly delayed growth and decreased survival after diamide and quinone stress, but no growth phenotype under sublethal oxidative stress as well as antibiotics treatment

To study the function of Ap_4_A in *S. aureus*, we performed growth phenotype analyses of the Δ*yqeK* mutant and the *yqeK*^+^ complemented strain. The Δ*yqeK* mutant was significantly impaired in growth in Luria broth (LB), Tryptic soy broth (TSB) and RPMI medium and reached the stationary phase about 1-2 h later than the WT (**Fig. 2A-C**). The growth was restored to WT levels in the *yqeK*^+^ complemented strain in TSB and RPMI medium, indicating that the increased levels of AP4N cause a fitness defect in *S. aureus*. In LB medium, ectopic expression of *yqeK* partially restored the growth defect of the Δ*yqeK* mutant to WT level (**Fig. 2A**). To analyze the phenotypes after exposure to oxidative and electrophile stress, the *S. aureus* strains were grown in RPMI to the exponential growth phase at OD_500_ of 0.5 and treated with sublethal concentrations of 50 µM methylhydroquinone (MHQ), 1.75 mM hypochlorous acid (HOCl), 10 mM hydrogen peroxide (H_2_O_2_) or 2 mM diamide to monitor the growth (**Fig. S1A-H**). However, the Δ*yqeK* mutant and the *yqeK*^+^ complemented strains did not show significantly increased susceptibilities *versus* the WT after exposure to MHQ, HOCl, H_2_O_2_ and diamide stress (**Fig. S1A-H**), indicating that increased Ap_4_A levels do not impact the growth during acute sublethal oxidative or electrophile stress in *S. aureus*. To analyze whether the YqeK affects the survival after exposure to lethal oxidative and electrophile stress, we treated the *S. aureus* strains with 5 mM diamide and 100 µM MHQ and determined the survival rate after 2 and 4 hours by CFU counts (**Fig. 2D, E**). While the survival of the Δ*yqeK* mutant and *yqeK*^+^ complemented strains was not affected after 2 hours, the viability of the Δ*yqeK* mutant was significantly 2- to 3-fold decreased after 4 hours of diamide and MHQ stress and restored to WT level in the and *yqeK*^+^ strain (**Fig. 2D, E**). These results indicate that YqeK does not play an important role during acute sublethal stress, but rather improves the survival after long-term exposure to toxic oxidative and electrophile stress. Additionally, we investigated the antibiotic tolerance of the *S. aureus* COL Δ*yqeK* mutant and *yqeK*^+^ complemented strains after exposure to sublethal doses of gentamicin, rifampicin and ciprofloxacin (**Fig. 3A-F**). However, the growth comparison did not show any significant difference between the antibiotic-treated Δ*yqeK* mutant or *yqeK*^+^ complemented strains *vs*. the WT. Thus, Ap_4_A does not contribute to antibiotics resistance in growth experiments in *S. aureus*.

**Fig. 2.**
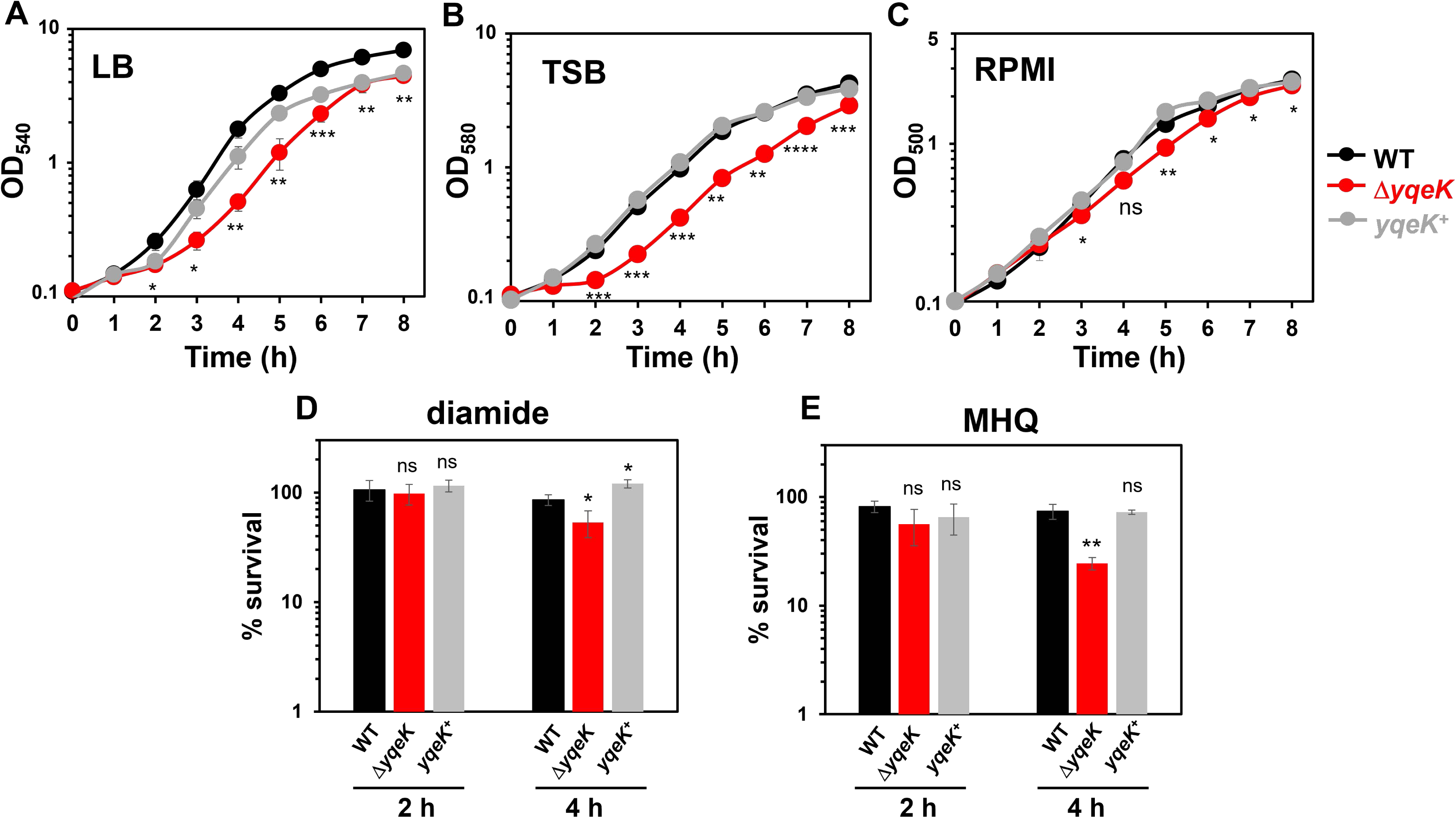
The Δ*yqeK* mutant shows a growth delay in TSB, LB, and RPMI medium and decreased survival after diamide and quinone stress. **(A-C)** The growth curves of *S. aureus* COL WT, the Δ*yqeK* mutant and *yqeK*^+^ complemented strains were monitored in LB **(A)**, TSB **(B)** and RPMI **(C)**. **(D-E)** For survival assays, *S. aureus* COL strains were grown in RPMI medium to an OD_500_ of 0.5 and exposed to lethal 5 mM diamide (Dia) **(D)**, or 100 µM methylhydroquinone (MHQ) stress (**E)**. Survival rates were determined after 2 and 4 hours relative to the untreated control, which was set to 100%. Results are shown as mean values of three independent biological experiments with error bars representing the SD. Statistical significance was calculated by comparing the Δ*yqeK* mutant *vs*. WT, and additionally *yqeK*^+^ *vs*. WT in Fig. 2 D, E using an unpaired two-tailed Student’s *t*-test (^ns^, *p* > 0.05; *, *p* ≤ 0.05; **, *p* ≤ 0.01; ***, *p* ≤ 0.001).

**Fig. 3.**
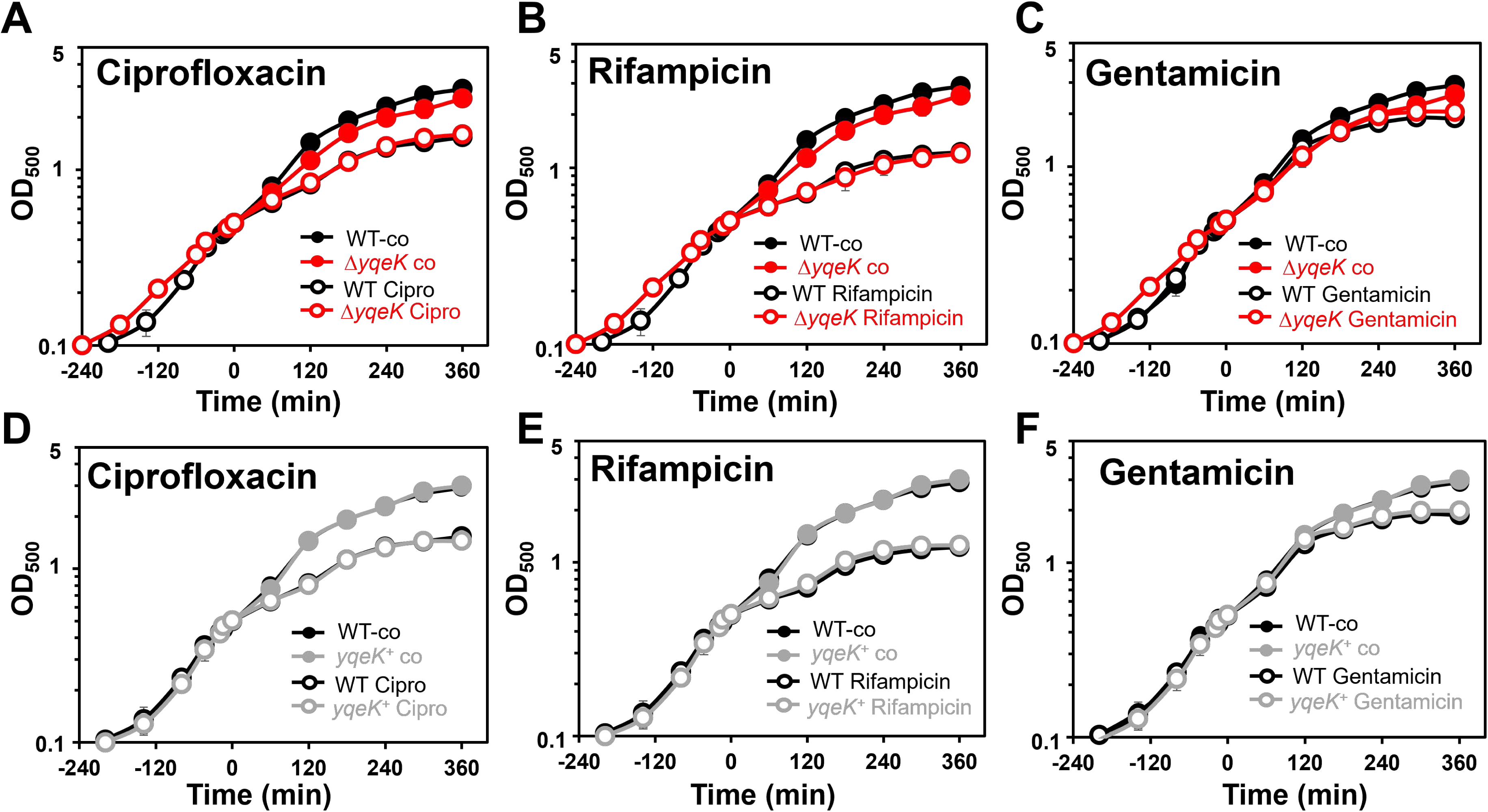
The Δ*yqeK* mutant does not show phenotypes after exposure to the antibiotics ciprofloxacin, rifampicin and gentamicin. The *S. aureus* COL WT, Δ*yqeK* mutant and *yqeK*^+^ complemented strains were grown in RPMI to an OD_500_ of 0.5 and treated with 90.5 µM ciprofloxacin **(A, D)**, 0.1 µM rifampicin **(B, E)**, or 0.1 µM gentamicin **(C, F)**. Data are presented as mean values of three biological replicates with error bars indicating the SD. No significant differences were calculated for the comparison of the Δ*yqeK* mutant or the *yqeK*^+^ complemented strain *vs*. the WT using an unpaired two-tailed Student’s *t*-test.

### RNA-seq transcriptomics reveals altered amino acid, nucleotide and iron metabolism and virulence gene expression in the *S. aureus* Δ*yqeK* mutant

Next, we performed RNA-seq transcriptome comparison of the *S. aureus* COL WT, the Δ*yqeK* mutant and the *yqeK^+^* complemented strain grown in TSB and harvested at an OD_580_ of 2. For significant fold-changes, we have chosen the M-value cut-off (log2-fold change) of ≥1 and ≤-1 for the comparison Δ*yqeK vs*. WT and *yqeK*^+^ *vs*. WT. Based on this cutoff, 123 genes were significantly >2-fold upregulated and 107 genes were <0.5-fold downregulated in the transcriptome of the Δ*yqeK* mutant *vs*. WT. The transcriptome comparison of *yqeK*^+^ *vs*. the WT resulted in 98 genes significantly >2-fold upregulated and 8 genes <0.5-fold downregulated transcripts (**Figs. 4, 5; Tables S1-S3**).

**Fig. 4.**
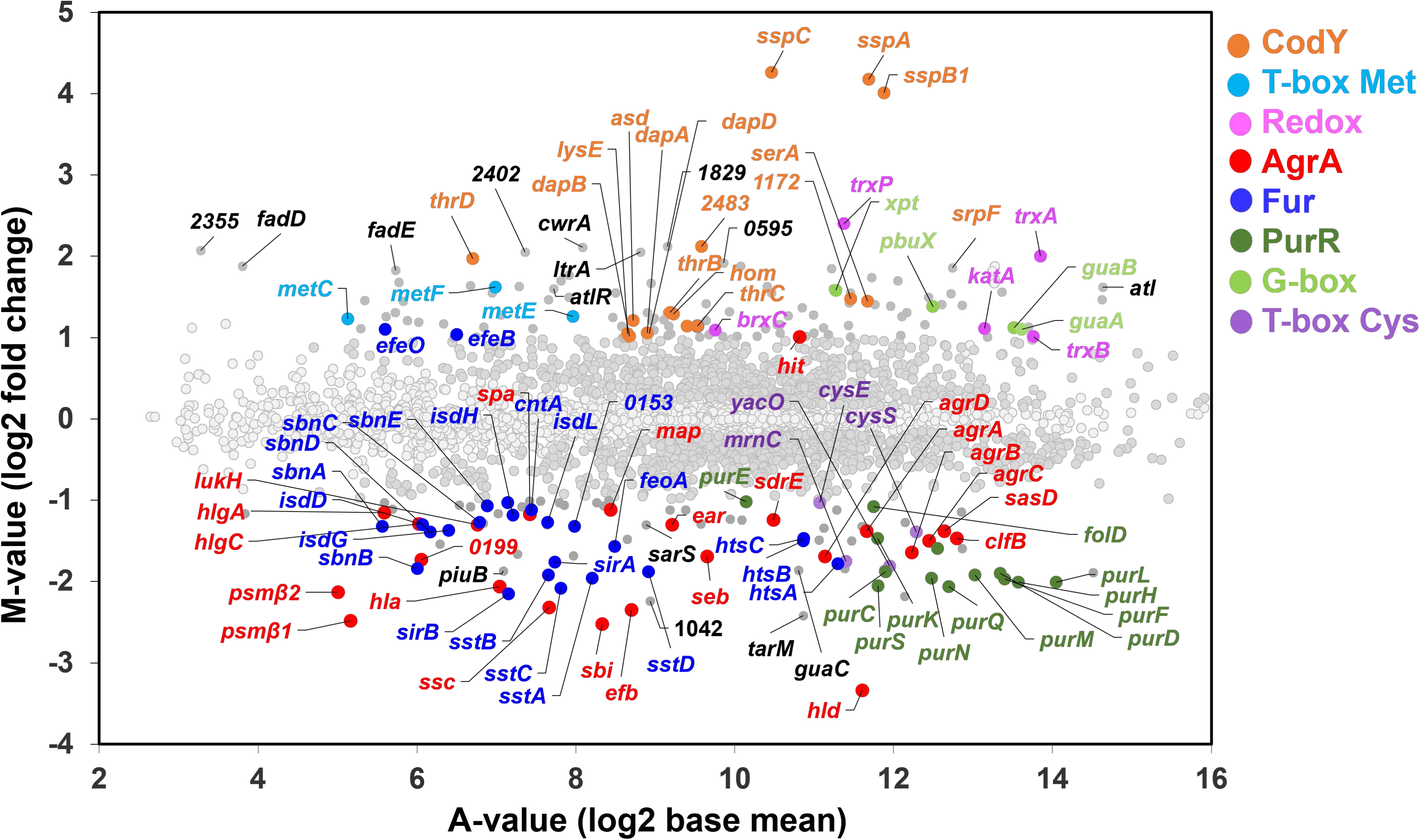
M/A-plot of the RNA-seq transcriptome results of the Δ*yqeK* mutant *vs*. the WT reveals induction of amino acid and GTP biosynthesis and downregulation of virulence factors, iron acquisition and purine biosynthesis. *S. aureus* COL WT and the Δ*yqeK* mutant were grown in TSB with 1% (w/v) glucose and 1% (w/v) xylose and harvested at an OD_580_ of 2 for RNA isolation. The gene expression profile of the Δ*yqeK* mutant *vs*. the WT is shown as a ratio/intensity scatterplot (M/A-plot), based on differential gene expression analysis using DESeq2 implemented in ReadXplorer v.2.2. Colored symbols indicate significantly induced or repressed transcripts (M-value ≥ 1 or ≤ −1; adjusted *p*-value ≤ 0.01), which were allocated to specific regulons or functional groups. Induced regulons include CodY (orange), redox-regulated genes (pink), T-box Met (cyan) and G-box (light green). Repressed regulons are AgrA (red), Fur (dark blue), T-box Cys (purple) and PurR (dark green). Light gray symbols denote transcripts below the significance cut-offs (adjusted *p*-value > 0.05 or absolute M-value < 1). The complete transcriptome data and regulon classifications of the datasets Δ*yqeK* mutant *vs*. WT and yqeK^+^ complemented strains *vs*. WT are presented in **Tables S1, S2, and S3**.

**Fig. 5.**
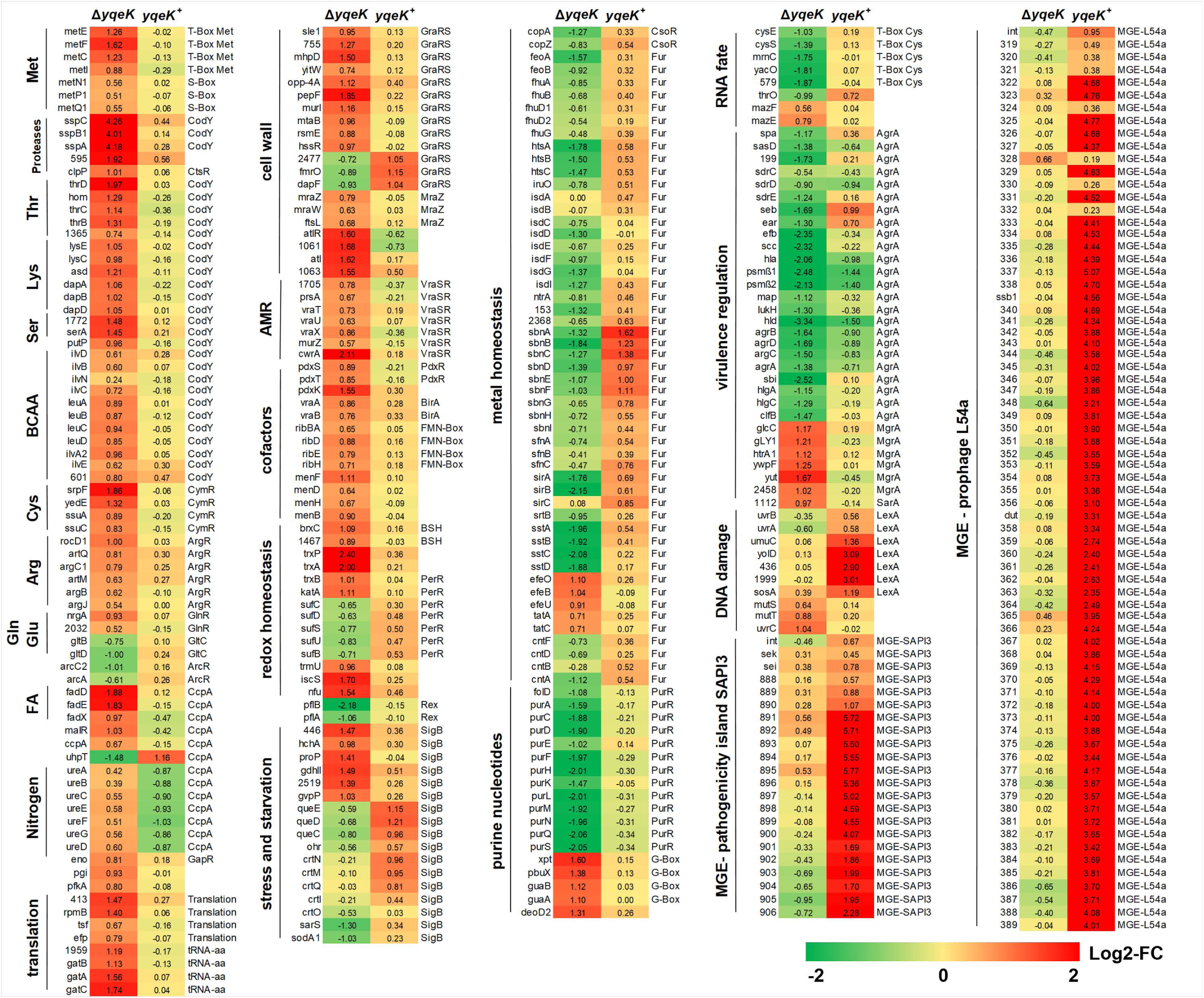
Heatmap for transcriptome comparison of the Δ*yqeK* mutant and *yqeK* complemented strains *vs*. the WT. *S. aureus* COL WT, Δ*yqeK* mutant and *yqeK*^+^ complemented strains were grown in TSB medium with 1% (w/v) glucose and 1% (w/v) xylose and harvested at an OD_580_ of 2 for RNA isolation. The heatmap shows differentially expressed regulons in the Δ*yqeK* mutant *vs*. WT and in the *yqeK*^+^ strain *vs*. the WT based on DESeq2 analyses integrated in ReadXplorer v2.2. The M-values (log2 fold-changes) of gene expression changes in the Δ*yqeK* mutant or the *yqeK*^+^ strain *vs*. the WT are shown using a red-yellow-green color code, where red indicted induction and green repression. The RNA-seq transcriptome datasets including detailed functional categories and regulon classifications are presented in **Tables S1-S3**.

The transcriptome results showed that the CodY-controlled *sspABC* operon encoding for extracellular serine and cysteine proteases and their inhibitors was most strongly (16 to 19-fold) upregulated in the Δ*yqeK* mutant (**Figs. 4, 5; Table S1-S3**). Additionally, other CodY regulon members were significantly 1.5 to 3.9-fold induced in the Δ*yqeK* mutant, including operons for the biosynthesis of threonine and homoserine (*hom-thrC-thrB*), lysine (*lysC-asd-dapA-dapB-dapD*), serine (*SACOL1772-serA*) and branched chain amino acids (*ilvDBNC-leuABCD-ilvA2*) that are upregulated due to GTP depletion upon AP4N accumulation (**Figs. 4, 5; Tables S1-S3**). Apart from the CodY regulon, amino acid biosynthesis genes for methionine, arginine and ornithine were 1.5 to 3-fold upregulated, including the T-box Met-regulated *metEFCI* operon and the ArgR-dependent *argBJCD* and *artQM* operons, whereas only few genes of the CymR and GlnR regulons for glutamine (*nrgA*) and cysteine biosynthesis (*ssuAC, srpF, yedE*), respectively, were 1.5 to 3.9-fold induced. Furthermore, the CcpA-controlled *fadDEX* operon for fatty acid degradation and the *ureABCEFGD* operon for urea degradation showed 1.5 to 3.7-fold increased expression in the Δ*yqeK* mutant (**Figs. 4, 5; Tables S1-S3**).

The transcriptome data further suggest alterations in cell wall biosynthesis and cell wall stress in the Δ*yqeK* mutant as revealed by the induction of some GraRS and VraRS regulon genes, e.g. *cwrA* (4.3-fold), the endopeptidase *pepF* (3.6-fold), the autolysin-encoding *atlR-1061-atl-1063* cluster (3-fold) and the MraZ-controlled division and cell wall (*dcw*) operon (1.6-fold) (**Figs. 4, 5; Tables S1-S3**). We further noted the 1.5- to 2.9-fold upregulation of the FMN-box, BirA and PdxR regulons for riboflavin, biotin, thiamine and menaquinone cofactor biosynthesis upon YqeK depletion. Additionally, the transcriptome results showed the 2- to 3-fold induction of the G-box regulon, including the *xpt-pbuX-guaB-guaA* operon, which is involved in GMP-biosynthesis. Moreover, the genes encoding the catalase *katA*, thioredoxins (*trxA, trxP*), thioredoxin reductase (*trxB*) and bacilliredoxin C (*brxC*) were 2-5-fold upregulated, suggesting increased ROS levels and an impaired thiol-redox homeostasis in the Δ*yqeK* mutant (**Figs. 4, 5; Tables S1-S3**). While transcription of *katA* and *trxB* is controlled by the PerR repressor (50), other PerR regulon members, such as the *ahpCF* peroxiredoxin, *dps* for iron storage and the *sufCDSUB* operon for FeS cluster biogenesis are 0.4 to 0.8-fold downregulated in the Δ*yqeK* mutant (**Tables S1-S3**). However, the *trmU-iscS* operon and *nfu* involved in FeS cluster biogenesis are 3-fold upregulated in the Δ*yqeK* mutant, perhaps as compensatory mechanism for *sufCDSUB* operon repression.

Furthermore, the Agr, Fur, PurR and T-box Cys regulons were significantly downregulated in the Δ*yqeK* mutant, indicating repression of Agr-controlled virulence factors, Fur-dependent iron-uptake systems, purine nucleotide biosynthesis and cysteinyl-tRNA-synthetase (**Figs. 4, 5; Tables S1-S3**). Specifically, the transcription of 24 genes of the Agr virulence regulon was 0.1 to 0.5-fold decreased in the Δ*yqeK* mutant, including the *agrABCD* operon, α, γ and δ−hemolysins (*hla, hlgAB, hld*), leukocidins (*lukH*), staphylocoocal enterotoxin B (*seb*), phenol soluble modulins (*psm*β*1* and *psm*β*2*), and cell surface-anchored adhesins (*spa, sbi, sdrCD, sdrE, efb, clfB, sasD*) (**Figs. 4, 5; Tables S1-S3**) (51–54). Additionally, 42 Fur regulon members were 0.2 to 0.7-fold downregulated in the Δ*yqeK* mutant, including the *sirABC, sbnABCDEFGHI, sfnABC, sstABCD isdCDEF-srtB-isdG, htsABC* and *SACOL2471-cntFDCBA* operons for the biosynthesis and transport of the siderophores staphyloferrin A and B, for the uptake of heme, catecholate and hydroxamate xenosiderophores and to transport the staphylopine metallophore to capture ferric iron (21, 55–57). However, the Fur-controlled *efeOBU* operon encoding a dye-decolorizing (DyP)-type heme peroxidase was 2-fold upregulated in the Δ*yqeK* mutant. Overall, the repression of the Fur regulon might indicate higher iron levels in the Δ*yqeK* mutant compared to the WT. Furthermore, consistent with the decreased levels of the adenylate and guanylate nucleotide pools due to accumulation of AP4N, transcription of the *purEKCSQLFMNHD* operon for purine biosynthesis was significantly (0.2 to 0.5-fold) downregulated in the Δ*yqeK* mutant (**Figs. 4, 5; Tables S1-S3**). Additionally, the T-box Cys-controlled *cysE-cysS-mrnC-yacO-SACOL0579* operon, encoding for an O-acetyl-cysteine synthase, a cysteinyl-tRNA-synthetase and a RNA methyltransferase (58), was 0.27 to 0.5-fold downregulated in the absence of YqeK. These results suggest the repression of tRNA^Cys^ loading with cysteine perhaps as consequence of increased thiol-oxidation and an impaired thiol-redox balance.

The transcriptome comparison of the *yqeK*^+^ complemented strain *vs*. the WT showed that the expression of the majority of up- and downregulated regulons in the Δ*yqeK* mutant was restored back to WT levels or even regulated opposite in the *yqeK*^+^ as compared to the Δ*yqeK* mutant. For example, the CcpA-dependent *ureABCEFGD* urease operon was 0.5-fold downregulated and the GraRS-regulated *SACOL2477-fmrO-dapF* operon and the Fur-controlled *sbnABCDEFGHI* operon for staphyloferrin A were 1.4-3-fold upregulated in the *yqeK^+^* strains, opposite to the Δ*yqeK* mutant *vs*. WT comparison, supporting that gene regulatory changes in the Δ*yqeK* mutant are related to the absence of YqeK and overproduction of Ap_4_A (**Fig. 5; Tables S1-S3**). However, expression of the strongly downregulated Agr regulon genes in the Δ*yqeK* mutant (e.g. *agrABCD*, *hld, hla, psmß1* and *psmß2* and *sdrCE*) could be only partially ∼50% restored to WT level in the *yqeK*^+^ strain, probably because Agr shows variations along the growth curve. Additionally, we noted that the *yqeK*^+^ complemented strain showed a DNA damage SOS response, as revealed by the induction of few LexA regulon members, including the *uvrAB* excinuclease operon (1.5-fold)*, umuC, yolD sosA, SACOL0436* and *SACOL1999* (2.3 to 8.5-fold) encoding small hypothetical proteins. Furthermore, transcription of large operons encoding MGE was strongly upregulated in the *yqeK*^+^ complemented strain, including the prophage L54a operon *SACOL0318-0389* (1.3 to 27-fold) and the 15.9 kb pathogenicity island SaPI3 operon *SACOL0885-0908* (1.5 to 55-fold) encoding staphylococcal enteroxins (*sek, sei, seb, ear*) (59), supporting the mobilization of MGEs by DNA damage due to overproduction of YqeK in *S. aureus* (**Fig. 5; Tables S1-S3**).

Overall, the gene expression profile of the Δ*yqeK* mutant indicates that enhanced Ap_4_A levels are associated with the upregulation of the biosynthesis of amino acids and GTP as well as oxidative stress, whereas the expression of virulence factors, iron uptake systems, purine biosynthesis and cysteinyl-tRNA synthetase are downregulated. These regulatory changes in the Δ*yqeK* mutant were successfully restored to WT level upon complementation, which also caused a DNA damage response resulting in induction of the prophage L54 and the pathogenicity island SaPI3.

### qRT-PCR confirms altered transcription of the *sspABC* operon, *agr* and *trxP* in the *S. aureus* Δ*yqeK* mutant

qRT-PCR analysis was used to validate the gene expression changes in the Δ*yqeK* mutant *vs*. the WT throughout the growth curve. Specifically, we analyzed transcription of the downregulated Agr regulon genes *agrA, RNAIII, hla* and *psmß1* at different time points (OD_540_ of 2, 3, and 4) along the growth in LB medium (**Fig. 6**). Consistent with the RNA-seq profile, transcription of *agrA, RNAIII, hla*, and *psmß1* was about 0.5-fold downregulated as compared to the WT and the complemented strain, indicating that expression of the Agr regulon is decreased due to the absence of YqeK. Similarly, transcription of the CodY-regulated *sspA* gene was strongly 3 to 6-fold induced along the growth in the absence of YqeK, and restored to WT level upon complementation, supporting the transcriptome data (**Fig. 6**). The qRT-PCR analyses further showed the 3 to 6-fold induction of the thioredoxin encoding *trxP* gene in the Δ*yqeK* mutant, whereas only significant changes of the SigmaB regulon gene *asp23* were observed for the OD 3 time point. Finally, the strongly enhanced *yqeK* transcription in the complemented strain could be quantified using qRT-PCR (**Fig. 6**). Overall, our qRT-PCR analyses validated the downregulation of the Agr virulence regulon and the strong induction of the *sspABC* operon and *trxP* gene in the absence of YqeK as revealed by the RNA-seq transcriptome data.

**Fig. 6.**
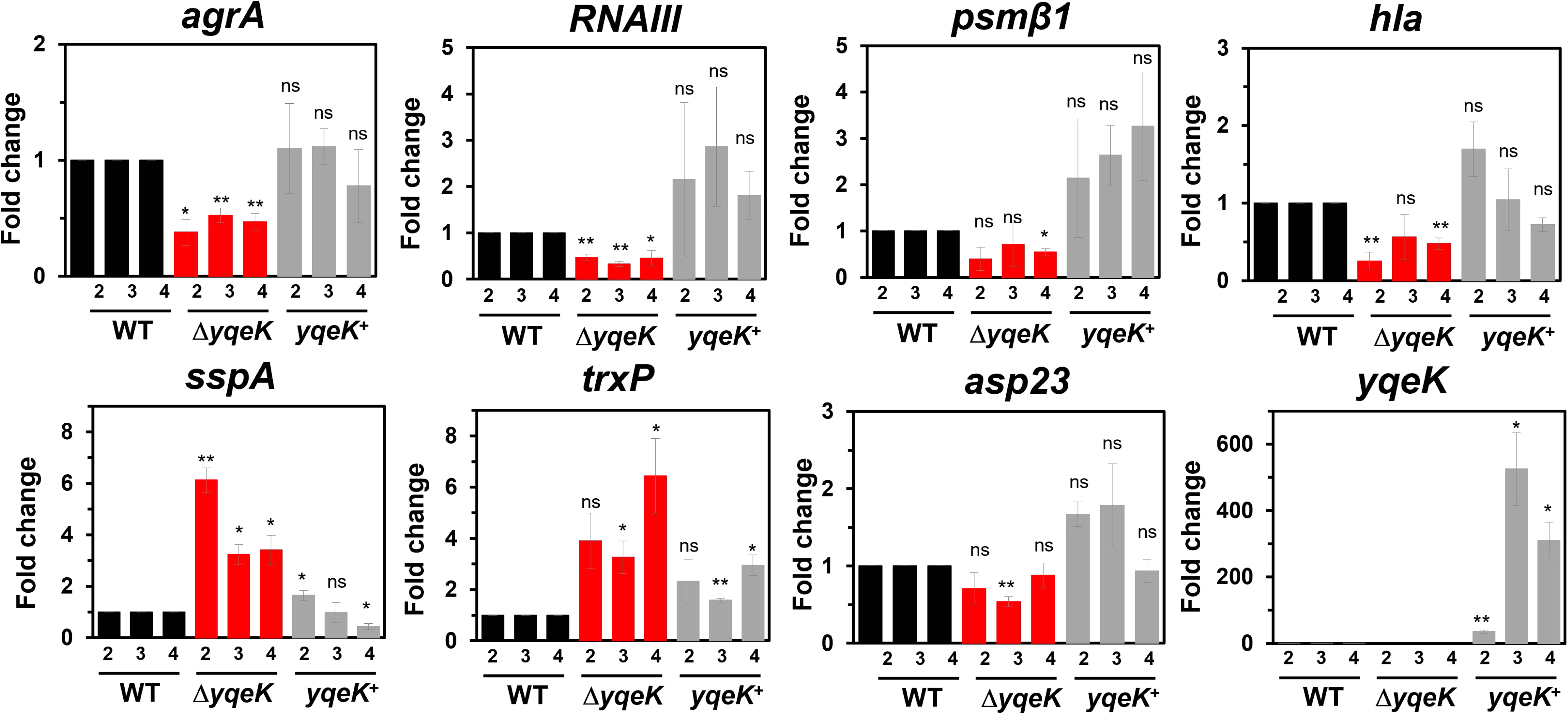
qRT-PCR analysis indicates downregulation of *agrA*, *RNAIII*, *psm*β*1* and *hla*, as well as increased transcription of *sspA* and *trxP* in the Δ*yqeK* mutant. *S. aureus* COL WT, the Δ*yqeK* mutant and the *yqeK*^+^ complemented strain were grown in LB with 1% (w/v) glucose and 1% (w/v) xylose in 3 biological replicates. Samples were harvested at OD_540_ of 2, 3 and 4. The relative transcript levels of *agrA*, *RNAIII*, *psm*β*1*, *hla*, *sspA*, *trxP*, *asp23*, and *yqeK* were determined by qRT-PCR. Equal amounts of total RNA (5 µg) were used for cDNA synthesis in a single batch reaction. The fold-changes were calculated as 2^−ΔCt relative to the WT at each OD, which was set to a fold-change of 1 (black bars). Error bars represent the standard deviation of 3 biological replicates, each cDNA sample was measured in four technical qRT-PCR replicates. Statistical significance was determined for the comparison of Δ*yqeK* mutant or *yqeK*^+^ complemented strains *vs*. WT using an unpaired two-tailed Student’s *t*-test (^ns^, *p* > 0.05; *, *p* ≤ 0.05; **, *p* ≤ 0.01).

### The Δ*yqeK* mutation has an impact on biofilm formation, staphyloxanthin production, iron levels, and survival in macrophage infection assays

We were interested whether the decreased expression of the Agr virulence regulon leads to changes in virulence phenotypes in the Δ*yqeK* mutant (**Fig. 7**). Biofilm formation was quantified using crystal violet staining of the *S. aureus* COL strains grown in TSB with 1% glucose (w/v) in microtiter plate wells for 24 and 48 hours. The results showed a two-fold increased biofilm formation of the Δ*yqeK* mutant compared to the WT and complemented strain (**Fig. 7A**). To investigate whether the increased biofilm formation of the Δ*yqeK* mutant is associated with the downregulation of the Agr-controlled hemolysins and PSMs, we analyzed biofilm formation of the *S. aureus* USA300 Δ*agr*, Δ*psm*α, Δ*psm*β, and Δ*psm*αβ mutants compared to their isogenic USA300 JE2 and USA300 LAC WT strains using crystal violet staining. In agreement with previous studies (60–63), the Δ*agr*, Δ*psm*β, and Δ*psm*αβ mutants showed significantly 1.5-fold increased biofilm formation, suggesting that the downregulation of Agr-controlled PSMβ toxins might contribute to increased biofilm formation in the absence of YqeK (**Fig. 7B**).

**Fig. 7.**
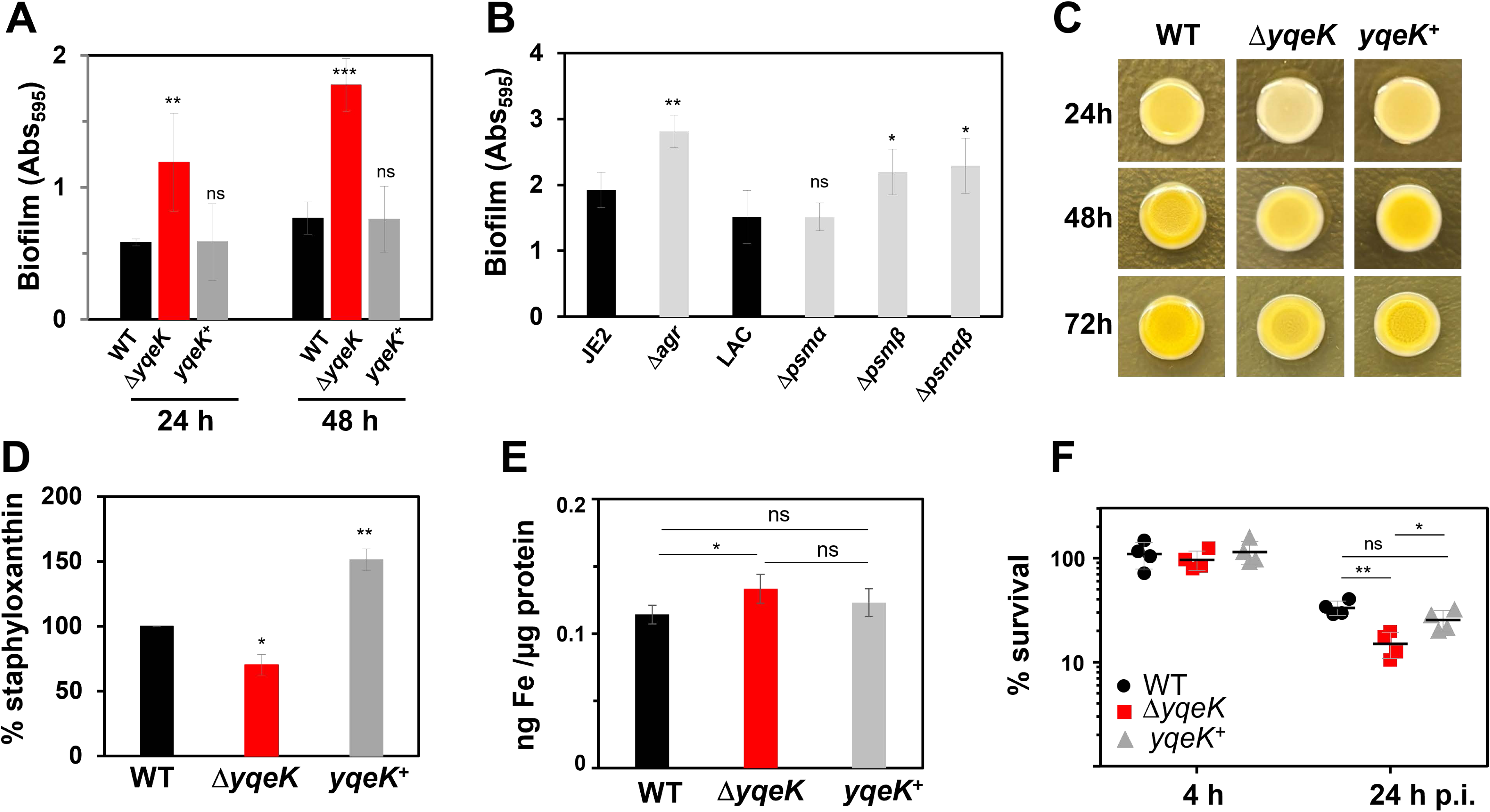
The Δ*yqeK* mutant shows increased biofilm formation, decreased staphyloxanthin pigmentation, enhanced intracellular iron level, and impaired survival during infection of murine J774A.1 macrophages. **(A)** *S. aureus* COL WT, Δ*yqeK* mutant and *yqeK*^+^ complemented strains were grown in TSB with 1% (w/v) glucose and 1% (w/v) xylose. Biofilm formation was analyzed using crystal violet staining from overnight cultures diluted to an OD_580_ of 0.5 in TSB with 1% (w/v) glucose and 1% (w/v) xylose. Cells were transferred in triplicate into sterile flat-bottomed 96-well microtiter plates and incubated at 37°C for 24 h and 48 h, followed by crystal violet staining. **(B)** Biofilm assays were performed using the *S. aureus* USA300 JE2 WT and the isogenic JE2 Δ*agr* mutant as well as the *S. aureus* USA300 LAC WT and the isogenic LAC Δ*psm*α, Δ*psm*β and Δ*psm*αβ mutants after 48 h using crystal violet staining. The results in A) and B) are average values of 3 independent biological experiments. Error bars represent the SD. Statistical significance of the results in A) and B) was determined using an unpaired two-tailed Student’s *t*-test (^ns^, *p* > 0.05; *, *p* ≤ 0.05; **, *p* ≤ 0.01; ***, *p* ≤ 0.001). **(C)** Macrocolony morphology and staphyloxanthin pigmentation were analyzed by spotting 2 µl of overnight cultures on TSB agar with 100 mM MgCl₂ followed by incubation for 24 h, 48 h and 72 h at 37°C. **(D)** Staphyloxanthin levels were quantified after 24 h of growth by methanol extraction from cell pellets of 1 ml cultures as described in the Methods section. Relative staphyloxanthin levels were calculated as 463/600 nm ratios and normalized to the WT, which was set to 100%. The results are shown as average values of 3 independent biological experiments. Error bars represent the SD. **(E)** Intracellular iron levels were quantified using the Ferene-s assay in the *S. aureus* COL WT, Δ*yqeK* mutant and *yqeK*^+^ complemented strains. Mean values of five biological replicates are presented and error bars represent the SD. The statistics of the results in D) and E) were calculated using an unpaired two-tailed Student’s *t*-test (^ns^, *p* > 0.05; *, *p* ≤ 0.05; **, *p* ≤ 0.01). **(F)** For infection assays, the survival of *S. aureus* COL WT, Δ*yqeK* mutant and *yqeK*^+^ complemented strains was analyzed at 4 h and 24 h post-infection (p.i.) of murine macrophage J774A.1 at a multiplicity of infection (MOI) of 1:1. The CFUs were determined and the percentage of survival was calculated relative to the 2 h time point, which was set to 100%. Results of 4 biological replicates with 2 technical replicates each are presented as scatter dots. Error bars represent the SD. Statistical significance was determined using one-way ANOVA with Tukey’s multiple comparisons post hoc test (^ns^, *p* > 0.05; *, *p* ≤ 0.05; **, *p* ≤ 0.01).

Biofilm formation of the Δ*yqeK* mutant was also assessed using structured macrocolonies developed over 3 days on TSB agar plates with MgCl_2_. While no differences in the structure of the macrocolonies were observed, the Δ*yqeK* mutant appeared more light-yellow colored after 24h, indicating reduced levels of staphyloxanthin of the Δ*yqeK* mutant (**Fig. 7C**). Quantification of the staphyloxanthin amounts was performed from the bacteria when grown in liquid culture in TSB medium for 24h, revealing 30% decreased staphyloxanthin levels in the Δ*yqeK* mutant, whereas the *yqeK*^+^ complemented strain produced 150% staphyloxanthin in comparison to the WT (**Fig. 7D**). Thus, the Δ*yqeK* mutant promotes biofilm formation through downregulation of PSMβ1/2, and produces lower levels of the carotenoid pigment staphyloxanthin.

Our RNA-seq data revealed a stronger Fur regulon repression in the absence of YqeK, including operons for uptake systems of iron, siderophores and heme, indicating possibly higher intracellular iron levels (**Fig. 4, 5; Tables S1-S3**). Thus, we used the ferene-s assay to quantify total intracellular iron levels in the Δ*yqeK* mutant and *yqeK*^+^ complemented strain. Indeed, the absorbance of ferene-s-Fe^2+^ complex was slightly but significantly higher in the Δ*yqeK* mutant as compared to the WT, suggesting that enhanced Ap_4_A levels lead to slightly higher intracellular iron levels, thereby decreasing the expression of Fur-controlled iron uptake systems (**Fig. 7E**).

Furthermore, infection assays with murine macrophages J774A.1 were conducted using the Δ*yqeK* mutant and the *yqeK^+^* complemented strain to determine the survival after 2, 4 and 24 hours post infection (p.i.) as compared to the WT (**Fig. 7F**). The viability rate of the WT was calculated as 33 ± 4.5 %, whereas the Δ*yqeK* mutant survived to 15 ± 3.7 % and the complemented strain showed 25 ± 5 % survival after 24 h p.i. compared to the 2 h time point p.i. Thus, the results from CFUs counts of infected macrophages revealed that the Δ*yqeK* mutant was significantly impaired in survival after 24 hours p.i., but the survival could be restored to WT level in the *yqeK*^+^ complemented strain (**Fig. 7F**). Altogether, our phenotype analysis revealed that YqeK plays an important role for fitness, biofilm formation, staphyloxanthin production and the survival during oxidative stress and macrophage infections in *S. aureus*.

## DISCUSSION

In this study, we identify YqeK as a major determinant of dinucleoside polyphosphate and nucleotide homeostasis in *S. aureus* COL and link this metabolic function to fitness, biofilm formation, oxidative stress resistance and virulence-associated phenotypes. Deletion of *yqeK* caused a ∼105-fold accumulation of Ap_4_A, together with increased Ap_4_U and Ap_4_G and a broad reduction of adenylate, guanylate, uridylate, and cytidylate nucleotide pools. These metabolic changes were accompanied by delayed growth, differential gene expression, increased biofilm formation and intracellular iron levels, reduced staphyloxanthin production, and impaired survival of lethal oxidative and quinone stress and during infection of murine J774A.1 macrophages. Moreover, the transcriptome revealed significant induction of the CodY, G-box and T-box Met regulons as well as repression of the Agr, Fur, PurR, and T-box Cys regulons in the absence of YqeK. Together, our results support a model in which YqeK-dependent control of Ap_4_N levels connects nucleotide homeostasis with metabolic and virulence regulation in *S. aureus* COL.

The strong accumulation of Ap_4_A in the Δ*yqeK* mutant provides *in vivo* evidence that YqeK is a principal Ap_4_A hydrolase in *S. aureus* COL. This finding extends previous biochemical studies showing that purified YqeK preferentially hydrolyzes Ap_4_A into two molecules ADP and is consistent with Ap_4_N accumulation reported for Δ*yqeK* mutants of other *Firmicutes*, including *B. subtilis* and *Streptococcus mutans* (37, 45–47, 64). The much stronger increase in Ap_4_A than in Ap_4_U or Ap_4_G in our metabolomic analysis further agrees with the biochemical preference of YqeK for Ap_4_A (37, 47). Importantly, Ap_4_N accumulation was associated with a broad decrease in canonical nucleotide pools, including ATP, ADP, AMP, GTP, GMP, UTP, UDP, CTP, and CMP. Thus, loss of YqeK does not simply alter the abundance of Ap_4_A but perturbs cellular nucleotide homeostasis more generally. This nucleotide imbalance and Ap4A accumulation provide a plausible metabolic basis for the delayed growth of the Δ*yqeK* mutant under different cultivation conditions in *S. aureus* COL. Similarly, Ap4N accumulation affected the growth of the Δ*yqeK* mutants in *S. aureus* USA300 LAC (49) and *B. subtilis* (47), and in the *P. aeruginosa* Δ*apaH* mutant (42), supporting that the lack of Ap_4_A removal leads to fitness costs in different bacteria.

A particularly prominent consequence of YqeK depletion was the remodeling of GTP- and amino acid-responsive gene expression. The Δ*yqeK* mutant displayed reduced guanylate nucleotide pools together with induction of the G-box-regulated *xpt-pbuX-guaB-guaA* operon and CodY-dependent amino acid biosynthesis and the SspA/B extracellular proteases. Because GTP acts as an important metabolic input for CodY in *S. aureus* (18, 19, 65), these transcriptional changes are consistent with the measured depletion of the GTP pool. Ap_4_A has previously been shown to bind and inhibit the IMP dehydrogenase GuaB in *B. subtilis*, thereby regulating GTP biosynthesis (39). A similar mechanism could contribute to the guanylate depletion observed here, although direct interaction between Ap_4_A and *S. aureus* GuaB remains to be demonstrated. Identification of Ap_4_A-binding proteins in *S. aureus* will therefore be important for determining whether GuaB or other nucleotide-metabolic enzymes directly connect Ap_4_A accumulation to the observed transcriptional response.

The most relevant consequence of this metabolic remodelling for pathogenicity was the repression of the Agr virulence regulon. RNA-seq revealed decreased expression of the *agrABCD* operon together with Agr- and RNAIII-controlled toxins (*hla, hld, hlgAB, lukH, psm*β*1*, *psm*β*2*) and cell wall-anchored adhesins (*spa, sbi, sdrCD, sdrE, efb, clfB, sasD*) (51–54). The qRT-PCR analyses confirmed the repression of some Agr-regulated genes throughout the growth, supporting a robust relationship between YqeK depletion and downregulation of Agr-controlled virulence factors. Importantly, this transcriptome signature was accompanied by impaired survival of the Δ*yqeK* mutant inside J774A.1 murine macrophages. These results establish a functional link between YqeK-dependent nucleotide homeostasis and modulation of virulence traits in *S. aureus*. They are also consistent with the reduced viability of a *S. aureus* USA300 LAC *yqeK* transposon mutant in a murine skin abscess model (66), and with reduced pathogenicity of the *P. aeruginosa* Δ*apaH* mutant in plant, insect and mouse models (42). The *P. aeruginosa* Δ*apaH* mutant also downregulated secreted toxins, extracellular enzymes, and quorum-sensing molecules in the transcriptome (42), consistent with the repression of the quorum-sensing Agr system in Δ*yqeK* mutants of *S. aureus* COL and *S. aureus* USA300 LAC (49).

How Ap_4_N accumulation leads to Agr downregulation remains unresolved. A decreased GTP pool and CodY derepression provide one possible connection between nucleotide metabolism and virulence factor regulation as CodY negatively regulates the *agrABCD* operon (18, 19, 65, 67). However, Agr-dependent virulence factors are repressed in the Δ*yqeK* mutant and therefore CodY-independently regulated. Similarly, regulation of the most strongly upregulated *sspABC* operon encoding extracellular serine and cysteine proteases is unclear in the Δ*yqeK* mutant. Extracellular proteases are regulated by a complex and interactive regulatory network, involving primary (SarS, SarR, Rot, MgrA, CodY, SaeR, and SarA) and secondary regulatory factors (12, 15, 16, 68–72), but whether accessory virulence gene regulators are modulated by Ap_4_N levels remains an open question for future research. Overall, the gene regulatory changes suggest that Ap_4_N accumulation produces a broader regulatory state involving several interacting metabolic and virulence regulators. An additional possibility is regulation at the RNA level. YqeK was recently shown to control nucleoside tetraphosphate RNA capping in *Firmicutes*, and Np_4_-capped RNAs accumulate in a *B. subtilis* Δ*yqeK* mutant (48). Because RNA capping can alter RNA stability, including regulatory RNAs involved in virulence control, determining the Np_4_-capped transcriptome of *S. aureus* will be an important next step. RNA-capping using the metabolite nicotinamide adenine dinucleotide (NAD^+^) has been already shown for the Agr-controlled dual function RNAIII in *S. aureus*, resulting in changes of toxin secretion (73). At present, however, a direct contribution of Np_4_-capping to Agr or RNA III regulation remains to be established.

The increased biofilm formation of the Δ*yqeK* mutant provides a second phenotype consistent with reduced Agr activity. Agr promotes biofilm maturation and dispersal in part through production of phenol-soluble modulins (PSMs), which act as surfactant-like peptides that facilitate channel formation, biofilm detachment and disruption of matrix interaction (60–62, 74, 75). In our experiments, deletion of *agr* or *psm*β*1/2* increased biofilm formation, suggesting that reduced Agr-dependent PSMβ production contributes to the biofilm phenotype of the Δ*yqeK* mutant. Additionally, increased expression of the major autolysin Atl could promote initial attachment and contribute to the FnbB1/2-dependent biofilm formation in the absence of YqeK (76). Moreover, increased Ap_4_A levels have been previously associated with altered biofilm structures in *B. subtilis* (47, 77), *S. mutans* (64) and *Pseudomonas fluorescens* (78), indicating that Ap_4_N accumulation controls structured community behaviour in different bacteria.

YqeK depletion also affected iron homeostasis. The Fur regulon involved in iron-siderophore biosynthesis and uptake, heme acquisition, and metallophore transport was repressed in the Δ*yqeK* mutant of *S. aureus* COL and *S. aureus* USA300 LAC (49). The accompanying increase in total intracellular iron in our work provides a physiological explanation for this transcriptional signature, because elevated cellular iron would promote Fur repression of iron-acquisition genes. These findings reveal an additional connection between Ap_4_N/nucleotide metabolism and iron homeostasis central for *S. aureus* physiology and host-pathogen interactions (20, 21). Similarly, iron-acquisition systems were downregulated in the *P. aeruginosa* Δ*apaH* mutant, contributing to the reduced virulence (42). The mechanism linking Ap_4_N accumulation to increased intracellular iron is currently unknown and cannot be distinguished from secondary effects of the altered metabolic state. Nevertheless, the agreement between the Fur transcriptional signature and the independently measured iron phenotype argues that iron homeostasis is genuinely altered following loss of YqeK.

The Δ*yqeK* mutant additionally produced substantially lower staphyloxanthin amounts. Because staphyloxanthin protects *S. aureus* against oxidative damage and contributes to fitness during infection (79), its reduction could further contribute to impaired pathogenicity.

Furthermore, our phenotype assays revealed that the removal of Ap_4_A is important for survival after long-term exposure to thiol-specific oxidative and quinone stress, consistent with the sensitivity of the *S. aureus* USA300 LAC Δ*yqeK* mutant towards nitrosative and acid stress, which also alters the thiol-redox homeostasis of *S. aureus* (49). Additionally, the transcriptome results showed the upregulation of genes involved in the defense against oxidative stress, encoding the catalase, thioredoxins, thioredoxin reductases and bacilliredoxin (*katA*, *trxA, trxB*, *trxP, brxC)*, which were partly also upregulated in the Δ*yqeK* mutant of *S. aureus* USA300 LAC (49). The thiol-disulfide reducing systems maintain the cellular thiol-redox homoeostasis of *S. aureus* (80), suggesting increased protein thiol oxidation in the Δ*yqeK* mutant under non-stress conditions. This also explains the sensitivity of the Δ*yqeK* mutant to survive lethal diamide stress, which has been shown to cause strongly increased protein thiol-oxidation, including *S*-thiolations in the redox proteome of *S. aureus* (80–82).

The metabolic, transcriptional and phenotypic changes of the Δ*yqeK* mutant were restored towards WT levels upon *yqeK* complementation, supporting their dependence on YqeK. However, the *yqeK*^+^ complemented strain, which expresses YqeK from a xylose-inducible promoter, also displayed induction of few LexA-regulated genes and operons encoding prophage L54a, and SaPI3, indicating genotoxic stress and mobilization of MGEs. In mammalian cells, high Ap_4_A levels are generated as side-reaction of DNA ligase III by genotoxic compounds and in DNA repair mutants, leading to inhibition of DNA replication initiation to prevent replication of damaged DNA (83). The data therefore suggest that both excessive accumulation and removal of Ap_4_N may perturb cellular physiology by causing oxidative and genotoxic stress. Future studies should investigate the effect of different YqeK levels on prophage L54a induction, mutation and replication rates in *S. aureus*.

Together, our results establish YqeK as a central regulator of Ap_4_N homeostasis in *S. aureus* and reveal that disruption of this homeostasis has consequences extending beyond nucleotide metabolism. We propose that Ap_4_N accumulation and the associated depletion of canonical nucleotide pools generate a metabolic state characterized by altered GTP/CodY signaling, repression of Agr-dependent virulence and Fur-controlled iron acquisition, enhanced biofilm formation, reduced staphyloxanthin production, and impaired survival under oxidative and quinone stress and during infection of murine macrophages. Therefore, YqeK plays an important role in Ap_4_N homeostasis by coordinating metabolism with biofilm- and virulence-associated lifestyles to promote survival of *S. aureus* in immune cells and in mouse infection models (66). Defining the direct molecular targets of Ap_4_A, including potential protein-binding partners and Np_4_-capped regulatory RNAs, will be essential to establish the mechanisms connecting this unusual nucleotide metabolite to virulence regulation in *S. aureus*.

## Materials and Methods

### Bacterial strains and growth conditions

Bacterial strains, plasmids and primers used in this study are listed in **Tables S4 and S5**. *E. coli* strains were grown in LB medium for cloning and expression of plasmids. *S. aureus* COL strains were cultivated in LB, TSB, or in RPMI1640 supplemented with 7.5 µM FeCl_₂_ and 2.05 mM L-glutamine. The *yqeK^+^* complemented strain carrying the pRB473-*yqeK* plasmid was cultivated in the presence of 10 µg/ml chloramphenicol and 1% (w/v) xylose for induction. To analyze the growth under oxidative and electrophile stress, *S. aureus* COL WT, the Δ*yqeK* mutant and the *yqeK^+^*complemented strain were grown in RPMI and treated with 50 µM MHQ, 1.75 mM HOCl, 10 mM H_2_O_2_ or 2 mM diamide at an optical density at 500 nm (OD_500_) of 0.5 as described previously (84). For survival assays, *S. aureus* stains were grown in RPMI medium and exposed to 100 µM MHQ or 5 mM diamide at an OD_500_ of 0.5, followed by plating for colony forming units (CFU) after 2 and 4 hours on LB agar plates. After overnight incubation at 37°C of the LB plates, the survival rates were determined and normalized to the WT, set as 100%. The compounds diamide, MHQ, sodium hypochlorite and H_2_O_2_ (35% w/v) used for the stress experiments were purchased from Sigma-Aldrich.

### RNA isolation, library preparation, and next-generation cDNA sequencing

For the RNA-seq transcriptome analysis, *S. aureus* COL WT, the Δ*yqeK* mutant and the *yqeK*^+^ complemented strain were grown in TSB with 1% (w/v) xylose and 1% (w/v) glucose to an OD_580_ of 2. *S. aureus* cells were harvested and disrupted in 3 mM ethylenediaminetetraacetic acid (EDTA)/ 200 mM NaCl lysis buffer using a Precellys24 ribolyzer. RNA isolation was performed using the acid phenol extraction protocol as described (85). The RNA quality was checked by Trinean Xpose (Gentbrugge, Belgium) and the Agilent RNA Nano 6000 kit using an Agilent 2100 Bioanalyzer (Agilent Technologies, Böblingen, Germany). The Pan-Bacteria riboPOOL rRNA removal kit from siTOOLs Biotech was used to remove the rRNA. The TruSeq Stranded mRNA Library Prep Kit from Illumina was applied to prepare the cDNA libraries. The cDNAs were sequenced paired-end on an Illumina NextSeq 2000 system (San Diego, CA) using 2x 50 nt read length. The transcriptome sequencing raw data files are available in the GEO database under accession number GSE344458 for the *S. aureus* COL Δ*yqeK vs*. WT and *yqeK*^+^ vs WT datasets, respectively used in this work for comparison.

### Bioinformatics RNA-seq data analysis

The obtained sequencing data were trimmed using TRIMMOMATIC v0.39 (86) in PE mode with the following parameters: validatePairs and trimmers ILLUMINACLIP:TruSeq3-PE-2.fa:2:30:10, SLIDINGWINDOW:4:15, and MINLEN:25. The trimmed reads were mapped against the complete genome of *S. aureus* COL [CP000046 (chromosome) and CP000045 (plasmid pT181)] using BOWTIE2 (87) with parameters -X 600. Raw read counts were determined using FEATURECOUNTS v2.0 (88), with parameters -O, -M, -t gene, -g ID, -s 2, and -p. The raw count matrix was normalized and analyzed using DESeq2 v1.52.0 (89) and APEGLM v1.34.0 (90) in R v4.6.1. Both matrices (raw and normalized counts) are available via GEO accession number GSE344458.

Differential gene expression analysis of *S. aureus* COL WT *vs*. Δ*yqeK* and *yqeK*^+^ *vs*. WT including normalization was performed from 3 biological replicates using Bioconductor package DESeq2 (89) included in the ReadXplorer v2.2 software (91). The signal intensity value (A-value) was calculated by log2 base mean of normalized read counts and the signal intensity ratio (M-value) by log2 fold-change. The evaluation of the differential RNA-seq data was performed using an adjusted *p*-value cut-off of *p* ≤ 0.01 and a signal intensity ratio (M-value) cut-off of ≥1 or ≤-1. Genes within the range of the cut-offs for *p*-values and *M*-values were considered as significantly differentially expressed in the Δ*yqeK* mutant or the *yqeK*^+^ strain in comparison to the WT.

### Construction of the *S. aureus* COL Δ*yqeK* mutant and *yqeK*^+^ complemented strain

The *S. aureus* COL Δ*yqeK* (*SACOL1649*) mutant was constructed as a markerless deletion by allelic replacement using the pMAD shuttle vector as described previously (84, 92). Briefly, the approximately 550 bp up- and downstream regions of the *yqeK* were amplified using two primer pairs listed in **Table S5**. Subsequently, the PCR products were fused by overlap extension PCR and cloned into the plasmid pMAD. The final plasmids were electroporated into the intermediate *S. aureus* RN4220 strain, followed by phage transduction into *S. aureus* COL using phage 81 (93). The clean marker-less Δ*yqeK* deletion mutant was selected and confirmed by PCR and sequencing.

For complementation, the *yqeK* gene was amplified from the genome of *S. aureus* COL using primers listed in **Table S5** and subsequently cloned into the pRB473 plasmid under control of the xylose-inducible promoter into the *Bam*HI and *Kpn*I restriction sites (94). The recombinant plasmid was then transferred to the *S. aureus* Δ*yqeK* mutant via phage transduction as described previously (84).

### Quantitative real-time PCR (qRT-PCR) analysis

*S. aureus* COL WT, the Δ*yqeK* mutant and the *yqeK*^+^ complemented strain were cultivated in LB supplemented with 1% (w/v) xylose and 1% (w/v) glucose and harvested at an OD_540_ of 2, 3 and 4. Total RNA was isolated as described above. RNA (5 µg) was reverse-transcribed into cDNA using the RevertAid Reverse Transcriptase (Thermo Fisher Scientific) according to the recommendations of the manufacturer. qRT-PCR analysis was performed using Blue S’Green qPCR 2X Mix (Biozym) and an iQ5 cycler detection system (Bio-Rad), 10-fold diluted cDNA and primers listed in **Table S5** as previously described (95). The fold-change was calculated as 2^−ΔCt relative to the WT at each OD, where ΔCt = Ct_sample − Ct_WT. The wild type was set to a fold-change of 1. The qRT-PCR analyses were performed using cDNA synthesized from three biological RNA replicates, with four technical qPCR replicates for each cDNA sample. Statistical significance was determined using a two-tailed unpaired Student’s *t*-test.

### Preparation of intracellular metabolites

*S. aureus* COL WT, the Δ*yqeK* mutant and the *yqeK*^+^ complemented strain were grown in RPMI 1640 with 1% (w/v) glucose and 1% (w/v) xylose in three independent biological replicates with three technical replicates each. One ml of culture was collected at OD_500_ of 1, added to 1 ml 70% methanol (v/v) pre-cooled at −80 °C for 48h and immediately mixed by inversion. Samples were centrifuged at 14,000 rpm for 10 min at −10 °C, the supernatant was carefully removed, and cell pellets were stored at −80 °C. For metabolite extraction, frozen cell pellets were resuspended in 200 µl extraction solution consisting of LC-MS grade methanol and TE buffer (10 mM Tris, 1 mM EDTA, pH 7.0) at a 1:1 (v/v) ratio, followed by addition of 200 µl chloroform. All extraction solvents were pre-chilled at −20 °C for at least 12 h prior to use. Samples were vortexed briefly and incubated for 2 h at 4 °C with shaking at 500 rpm. Phase separation was achieved by centrifugation at 14,000 rpm for 10 min at −10 °C. The upper aqueous phase was carefully collected, filtered through a 0.22 µm filter and stored at −80 °C.

### Quantification of dinucleoside polyphosphates and nucleotides by LC–MS/MS

Dinucleoside polyphosphates and nucleotides were quantified by liquid chromatography-tandem mass spectrometry (LC-MS/MS). Chromatographic separation was performed using an Agilent Infinity II 1290 HPLC system equipped with a SeQuant ZIC-pHILIC column (150 × 2.1 mm, 5 µm particle size, PEEK-coated; Merck). The column was maintained at 40 °C and operated at a constant flow rate of 0.2 mL/min. Mobile phase A consisted of 10 mM ammonium acetate in water (pH 9.0) supplemented with 5 µM InfinityLab Deactivator Additive (Agilent Technologies), whereas mobile phase B consisted of 10 mM ammonium acetate (pH 9.0) in 90% acetonitrile. The injection volume was 2 µL.

The chromatographic gradient was as follows: 0 – 1 min, 75% B; 1 – 6 min, linear decrease from 75% to 40% B; 6 – 9 min, 40% B; 9 – 9.1 min, linear increase from 40% to 75% B; and 9.1 – 20 min, 75% B for column re-equilibration.

Mass spectrometric analysis was performed using an Agilent 6470A triple quadrupole mass spectrometer equipped with an electrospray ionization source operated in negative ionization mode. The electrospray voltage was set to 3000 V and the nozzle voltage to 1000 V. The sheath gas temperature was 300 °C with a flow rate of 11 L/min, the nebulizer pressure was 20 psig, and the drying gas temperature was 100 °C with a flow rate of 11 L/min.

Target analytes were detected by multiple reaction monitoring (MRM) and identified based on their characteristic precursor-to-product ion transitions and retention times relative to authentic standards. Mass transitions, collision energies, cell accelerator voltages, and dwell times were individually optimized using authentic standards. Compound-specific MS parameters are provided in **Table S6**.

Chromatograms were integrated using MassHunter software (Agilent Technologies, Santa Clara, CA, USA). Analyte concentrations were quantified using external calibration curves generated from authentic standards covering a concentration range of 0 – 50 µM. Measured analyte concentrations were converted to absolute metabolite amounts using the extraction volume and subsequently normalized to biomass based on OD_500_ and culture volume. Normalized metabolite amounts are reported as nmol per OD500 x ml culture. The original metabolite data and normalized amounts are presented in **Table S7**.

### Biofilm formation, cultivation of macrocolonies and staphyloxanthin quantification

Biofilm formation was quantified using crystal violet staining in 96-well microtiter plates as described previously (96). Briefly, *S. aureus* strains were grown in TSB medium with 1% (w/v) glucose and 1% (w/v) xylose at 37 °C with a starting OD_580_ of 0.5 in a total volume of 200 µl. After 24 h and 48 h, biofilms were washed twice with 0.9% NaCl, stained with 0.1% crystal violet and disrupted with 1% SDS. Biofilm formation was quantified by measuring the absorbance at 595 nm.

*S. aureus* COL macrocolonies were grown on TSB agar supplemented with 100 mM MgCl_2_ as described previously (96). Briefly, 2 µl aliquots of overnight cultures were spotted onto plates and incubated for 3 days. Staphyloxanthin extraction was performed from 1 ml overnight cultures grown in TSB medium as described previously (96). Cell pellets were extracted three times in 250 µl methanol at 14,000 rpm and 55 °C for 3 min. Absorbance was measured at 463 nm using a CLARIOstar microplate reader (BMG Labtech) and staphyloxanthin levels were normalized to OD_600_ and expressed as A_463_/OD_600_ ratios.

### Ferene-s assay for quantification of intracellular iron level

The intracellular iron concentrations were determined using the ferene-s (3-(2-pyridyl)-5,6-di (2-furyl)-1,2,4-triazine-5′,5″-disulfonic acid disodium salt) (Sigma-Aldrich) as described previously with some modifications (27). Specifically, the *S. aureus* COL WT, Δ*yqeK* mutant and *yqeK*^+^ complemented strains were grown in TSB with 1% (w/v) xylose and 1% (w/v) glucose and harvested at an OD_580_ of 2. Cell pellets were lysed with 1% hydrochloric acid (HCl) and heated at 80 °C for 10 min. Excess acid was neutralized with 7.5% ammonium acetate. Ferric iron (Fe^3+^) was reduced to ferrous iron (Fe^2+^) using 4% ascorbic acid. The precipitated protein was complexed with 2.5% sodium dodecyl sulfate, followed by addition of 1.5% of the iron chelator ferene-s leading to the formation of the blue iron-ferene-s complex (Fe^2+^: ferene-s). Samples were centrifuged at 9000 rpm for 7 min and the absorbance was measured at 593 nm. Ammonium iron (II) sulfate hexahydrate (Sigma Aldrich) was used to prepare the iron standard curve.

### Infection assays with the murine macrophage cell line J774A.1

Intracellular survival assays were performed using the murine macrophage cell line J774A.1 as described previously (84). *S. aureus* COL WT, the Δ*yqeK* mutant and the *yqeK*^+^ complemented strain were used to infect macrophages at a multiplicity of infection (MOI) of 1. Extracellular bacteria were eliminated by treatment with 150 µg/ml gentamicin for 1 h. Intracellular survival of phagocytosed *S. aureus* was determined at 2, 4 and 24 hours post-infection by lysis of infected macrophages with 0.1% Triton X-100 and determination of CFUs on agar plates.

## ACKNOWLEDGEMENTS

This work was supported by a grant from the Deutsche Forschungsgemeinschaft, Germany (AN746/8-1) to H.A. G. B. acknowledges funding from the Deutsche Forschungsgemeinschaft within the GRK 2573 “Microbial Nucleotide Metabolism”. G.B. thanks the excellence cluster Microbes for Climate (M4C) for support. We would like to thank Christiane Wolz (Tübingen) for providing the *S. aureus agr*, *psm*α, *psm*β and *psm*αβ mutants. We further acknowledge Franziska Kiele for her excellent technical assistance in the survival assays.

## AUTHOR DISCLOSURE STATEMENT

No competing financial interests exist.

## DAT A AVAILABILITY STATEMENT

The authors confirm that the data supporting the findings of this study are available within the article and its supplementary materials. The transcriptome sequencing raw data files are available in the GEO database under accession number GSE344458.

