## Supplemental Figure 1 for "The diadenosine tetraphosphate hydrolase YqeK controls fitness, biofilm formation, staphyloxanthin production and virulence in *Staphylococcus aureus*"

Fig. S1

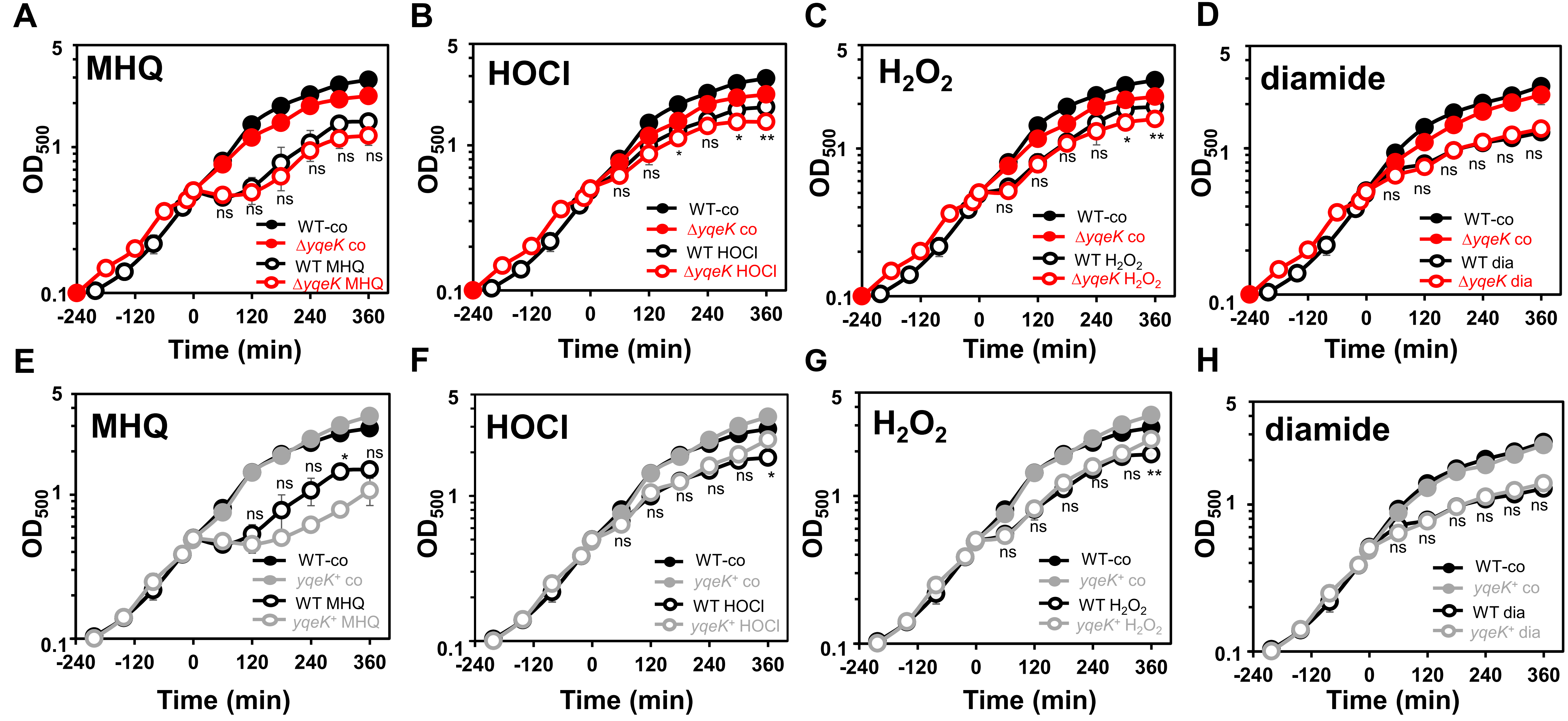


**Fig. S1. The Δ*yqeK* mutant is not impaired in growth after acute sublethal oxidative and electrophile stress.** *S. aureus* COL WT, Δ*yqeK* mutant **(A-D)** and *yqeK*⁺ complemented strains **(E-H)** were grown in RPMI with 1% glucose and 1% xylose to an $OD_{500}$ of 0.5, followed by exposure to 50 µM MHQ **(A, E)**, 1.75 mM HOCl **(B, F)**, 10 mM $H_2O_2$ **(C, G)**, or 2 mM diamide **(D, H)**. Data are shown as mean values of three biological replicates with error bars indicating the SD. Statistical significance was determined by comparison of the Δ*yqeK* mutant or the *yqeK*⁺ complemented strain *vs*. WT using an unpaired two-tailed Student's *t*-test WT ($^{ns}$, $p > 0.05$; *, $p \leq 0.05$; **, $p \leq 0.01$; ***, $p \leq 0.001$).
