## Supplemental Tables S4-S5 for "The diadenosine tetraphosphate hydrolase YqeK controls fitness, biofilm formation, staphyloxanthin production and virulence in *Staphylococcus aureus*"

**Table S4. Bacterial strains, phage and plasmids.**

| Strain | Description | Reference |
|---|---|---|
| ***Staphylococcus aureus*** | | |
| RN4220 | restriction negative strain/MSSA cloning intermediate derived from 8325-4 | (1) |
| COL | archaic HA-MRSA strain | (2) |
| COL-Δ*yqeK* | COL Δ*yqeK* deletion mutant | This study |
| COL-Δ*yqeK yqeK*$^+$ | COL Δ*yqeK* complemented with pRB473-*yqeK*$^+$ | This study |
| USA300 LAC | CA-MRSA strain | (3) |
| USA300 LAC Δ*psmα* | USA300 LAC Δ*psmα1,4* deletion | (4, 5) |
| USA300 LAC Δ*psmβ* | USA300 LAC Δ*psmβ1,2* deletion | (4, 5) |
| USA300 LAC Δ*psmαβ* | USA300 LAC Δ*psmα1,4*, Δ*psmβ1,2* deletion | (4, 6) |
| USA300 JE2 | CA-MRSA strain with cured plasmids | (7) |
| USA300 JE2 Δ*agr* | USA300 JE2 Δ*agr* deletion | (8) |
| *Staphylococcus* phage 81 | | (9) |
| **Plasmids** | | |
| pMAD | Temperature-sensitive *E. coli*/Gram-positive shuttle vector for allelic replacement and markerless gene deletion; Amp$^R$, Em$^R$ | (10) |
| pMAD-Δ*yqeK* | pMAD derivative carrying *yqeK* upstream and downstream flanking regions for Δ*yqeK* mutant construction | This study |
| pRB473 | pRB373-derivative, *E. coli*/ *S. aureus* shuttle vector, containing xylose-inducible $P_{Xyl}$ promoter Amp$^R$, Cm$^R$ | (11, 12) |
| pRB473-*yqeK* | pRB473 expressing *yqeK* under $P_{Xyl}$ | This study |

$^R$: resistant, Amp: ampicillin, Cm: chloramphenicol, Em: Erythromycin

**Table S5. Oligonucleotide (primer) sequences**

| Primer name | Sequence (5' to 3') |
|---|---|
| **Construction of the *S. aureus* COL Δ*yqeK* mutant** | |
| pMAD-yqeK-for-BglII | GTTACACATTAACTAGAC**AGATCT**CGTTCAGCACAGATTAACAATG |
| pMAD-yqeK-f1-rev | TTGTATGCCATATCTCGAATATCGAAATCATGTAATACACCTGCTA |
| pMAD-yqeK-f2-for | TAGCAGGTGTATTACATGATTTCGATATTCGAGATATGGCATACAA |
| pMAD-*yqeK*-rev-SalI | GATATCGGATCCATATGAC**GTCGAC**TTAATACGCAACCTGACTATAT G |
| **Construction of the *S. aureus* COL *yqeK*+ complemented strain** | |
| pRB-yqeK-for-BamHI | TAACCAACTAAAATGTA**GGATCC**ATATTAAGGGGGAAGGATTATATG |
| pRB-yqeK-rev-KpnI | CAGCGGAATTCGAGCTC**GGTACC**TTAATCATCCTTTATTCTTTCGTC |
| **qRT-PCR primers** | |
| qPCR-agrA-for | CGAGTCACAGTGAACTTACCTATT |
| qPCR-agrA-rev | AGCGTGTATGTGCAGTTTCTA |
| qPCR-RNAIII-for | AAACGACTGATGTGATGGAAA |
| qPCR-RNAIII-rev | AGTCACCGATTGTTGAAATGA |
| qPCR-psmβ1-for | CAGCAATTAATAATGATGGCGCAAA |
| qPCR-psmβ1-rev | CCGAATAATTTACCTAATAAACCTACGC |
| qPCR-hla-for | GAACCCGGTATATGGCAATCA |
| qPCR-hla-rev | GAGAACTTGCTTTGTTAGGATCAAG |
| qPCR-sspA-for | ATCGTCACCAAATCACAGATACA |
| qPCR-sspA-rev | ACAACTACACCGGAAGCAATAA |
| qPCR-trxP-for | CAGACTGTCGTGCTATGGATT |
| qPCR-trxP-rev | GCTAGGGATACCCATAACTTCATT |
| qPCR-yqeK-for | GAAATATTGCATGGCCCTGTG |
| qPCR-yqeK-rev | GTTGACGTCCAGTAGTATGGTATT |
| qPCR-asp23-for | GACTTCGATGATGGTCATGTTTAC |
| qPCR-asp23-rev | CGCAACTACATTTGGTTGTCTTA |

Restriction sites are bold
